# Choosing the best motion artefact correction: simplified and advanced how-to guides using QT-NIRS

**DOI:** 10.64898/2026.01.30.702862

**Authors:** Elizaveta Ivanova, Luca Pollonini, Mojtaba Soltanlou

**Affiliations:** University of Surrey, Faculty of Health and Medical Sciences, School of Psychology, UK; University of Houston, Department of Engineering Technology, Houston, TX, USA; University of Houston, Department of Electrical and Computer Engineering, Houston, TX, USA; University of Houston, Department of Biomedical Engineering, Houston, TX, USA; University College London, UCL Institute of Education, Department of Psychology and Human Development, London, UK; University of Johannesburg, Faculty of Education, NRF South Africa Chair in Integrated Studies of Learning Language, Science, and Mathematics in the Primary School, Johannesburg, South Africa

**Keywords:** fNIRS, QT-NIRS, motion artefacts correction, preprocessing, replicability

## Abstract

**Significance:** Selecting the appropriate motion artefact (MA) correction method for functional near-infrared spectroscopy (fNIRS) data is quite challenging, particularly in light of the need for standardised practice, replication, and transparency in the field. A clear framework for making measurable and replicable decisions is therefore essential.

**Aim:** This paper proposes a guide based on an open-source data quality assessment tool (QT-NIRS) that enables a transparent and evidence-driven choice of MA correction method.

**Approach:** We present the guide in two approaches: a simplified version that is easy to run for beginners, and an advanced version providing more informative output at the cost of additional computations and minor changes to the original QT-NIRS code. Due to its high flexibility and within-subject nature, the method is applicable across samples with varied characteristics.

**Results:** We applied the guide to two challenging datasets from 60 British preschoolers (mean age = 3.94 years, SD = 0.49) and 39 South African school children (mean age = 12.00 years, SD = 0.51). Both simplified and advanced approaches supported similar MA correction methods.

**Conclusions:** While both approaches can be used interchangeably, we recommend the advanced approach when possible due to its more informative and straightforward output, and advise caution when using the simplified version.

## 1 Introduction

Functional near-infrared spectroscopy (fNIRS) is an on-the-rise neuroimaging technique that uses near-infrared light to measure changes in oxygenated and deoxygenated haemoglobin as an indirect indicator of brain activity. A large advantage of fNIRS when compared to other neuroimaging techniques (e.g., EEG and fMRI) is that fNIRS can shed light–both literally and figuratively–on previously understudied populations, including infants (Bulgarelli et al., 2018; De Roever et al., 2018) and preschoolers (Ivanova et al., IPA), those in less-resourced (Fonseca et al., preprint) and remote geographical areas (Jasinska et al., 2024; Lloyd Fox et al., 2014), and in-bed patients (Casetta et al., 2024; Cutini & Brigadoi, 2014), due to its several strengths. Some of these strengths include tolerance for participant movement, high portability (some devices can easily fit in a backpack), wireless capability, and quick and easy setup (Soltanlou et al., 2018; Barreto & Soltanlou, 2022; Pinti et al., 2020). Although fNIRS has certain limitations, such as limited depth penetration, lower spatial resolution compared to fMRI, lower temporal resolution compared to EEG, and sensitivity to physiological noise (Providência & Margolis, 2022; Tachtsidis & Scholkmann, 2016), it remains an attractive option for studies involving populations that are typically difficult to test, such as young children, or for research that requires specific conditions, like tasks involving physical movement or naturalistic settings. As with any emerging technique, best practices for using fNIRS are still evolving (Yücel et al., 2021). Some of the recent attempts in society include standardising the best practices for the use of related terminology (Stute et al., 2025), planning experimental design (Schroeder et al., 2023), and publishing (Yücel et al., 2021) fNIRS studies.

An outstanding challenge in fNIRS research is the lack of consensus on preprocessing procedures (Yücel et al., 2025). Many researchers rely on their own established pipelines and often provide limited justification for their preprocessing choices. Yet, these steps can significantly influence results, as shown in the large-scale collaboration, known as the FRESH study by Yücel et al. (2025), in which 38 research groups preprocessed the same dataset using their own pipelines for analysis and came to a variety of different conclusions. Naturally, such alarming variability in results and conclusions confirms previous notions of the lack of reproducibility and replicability in fNIRS research (Kelsey et al., 2023; Schroeder et al., 2023; Yücel et al., 2021). With this in mind, continued dialogue and collaboration among researchers working with fNIRS to promote the development of standards for the analysis pipelines remains crucially important.

One crucial step of preprocessing–where the decisions are rather arbitrary and sometimes biased to the output–is the selection of the most suitable motion artefact (MA) correction method for the data at hand. The need comes from a two-fold problem. The first issue is that although several papers have measured the efficacy of various MA correction methods (e.g., Brigadoi et al., 2014; Fishburn et al., 2019; Gemignani & Gervain, 2021), the efficacy of these MA correction methods may differ across datasets. This directly relates to the second issue, which is the high variability of group characteristics, both physiological and task-related differences. For example, infants’ physiological differences often lead to cases when deoxygenated haemoglobin (HbR) responses increase together with oxygenated haemoglobin (HbO) responses, which is considered as noise in older children and adults (De Roever et al., 2018). Characteristics can also vary by task, depending on whether it is more or less engaging, which would then affect the number of MAs (e.g., an active response task would prompt participants to move more than a passive task, and more movements mean more MAs). Moreover, differences in skin and hair thickness can impact data quality, as darker skin can reduce the amount of light absorbed by the scalp, while thicker hair can prevent the optodes from achieving a consistently stable coupling with the scalp during recording (Bronkhorst et al., 2019; Kwasa et al., 2023). The existence of those group-and task-related differences means that even though some MA correction methods have been preferred for specific groups, such as Temporal Derivative Distribution Repair (TDDR) for school-age children (Fishburn et al., 2021) and Wavelet for infants (Gemignani & Gervain, 2021), the fit of these methods to certain groups is not guaranteed. Therefore, the challenge of developing a guideline for choosing the most suitable MA correction method for a data in hand remains unresolved.

In this paper, we propose an accessible data-driven approach for choosing the MA correction method using QT-NIRS–a widely used tool for quality check in fNIRS research (Hernandez & Pollonini, 2020). QT-NIRS splits the data into time windows (usually 3-5 seconds long) and assesses the quality of the data in each window based on the clear presence of heartbeat throughout the recording, relying on the scalp coupling index (SCI) that indicates cross-correlation between the signal’s wavelengths. As the optode movement during the recording can bias SCI, the peak spectral power (PSP) of the cross-correlated signal is used as a measure of the motion artefacts. When the number of bad windows (i.e., poor SCI and/or PSP) is higher than a certain percentage of the Single-Window Quality Threshold (QT, which is set by the user), such a time window is marked as excessively noisy. As the MA correction is supposed to improve the quality of data by reducing the number of bad windows, we suggest that by running QT-NIRS on the data pre-and post-MA correction, one can make an informed decision on which MA correction method is better by comparing the resulting metrics. To demonstrate this, we will apply the following MA correction to the two distinct datasets using these common methods (Santosa et al., 2020; Yucel et al., 2025): Principal Component Analysis (PCA; Zhang et al., 2005), Baseline PCA (Franceschini et al., 2006), Wavelet (Molavi & Dumont, 2012) and TDDR (Fishburn et al., 2021). Wavelet and TDDR were tested both individually and in combination as suggested by Guan et al. (2024).

Our proposed approach has three main goals. First, it will make the choice of the MA correction objective informed and replicable. Second, it will offer two options, simplified and advanced, to make it suitable for researchers with varying levels of programming skills. Third, it will be applicable across a wide range of datasets, not just for a certain population.

## 2 Methods

### 2.1 Datasets description and preprocessing

To demonstrate how our approach can work across different datasets, we applied it to two challenging cases. One dataset included preschool children in the UK, known for being very active, even when asked to sit still. The other dataset included primary school children in South Africa with characteristics that are known to present technical challenges for optical imaging, such as darker skin and thicker hair.

#### 2.1.1 Dataset A

Dataset A consisted of data from 60 neurotypical preschool-aged children (mean age = 3.94 years, SD = 0.49) residing in the UK. Haemodynamic responses were recorded during an approximately 9-minute-long active numerical task requiring verbal responses. Children were encouraged to remain still throughout the session, though age-related movement was expected during the data collection. A portable, continuous-wave fNIRS device (Brite, Artinis Medical Systems BV, The Netherlands) was used, equipped with 8 sources and 10 detectors emitting dual-wavelength near-infrared light (760 nm and 850 nm). Data were sampled at 25 Hz across 18 channels positioned bilaterally over the prefrontal and parietal cortices (Fig. 1).

**Fig. 1.**
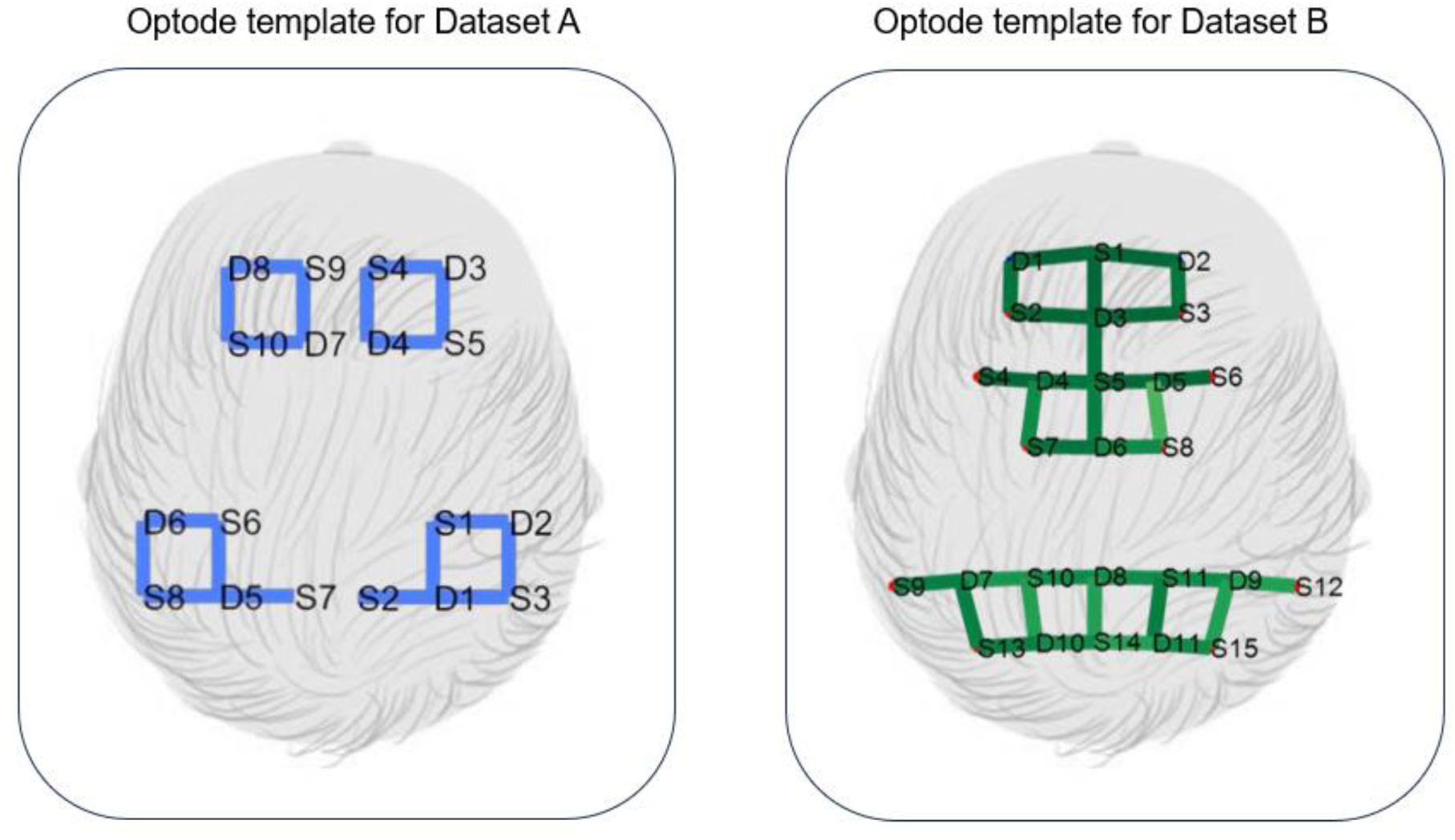
Optode templates for Dataset A (on the left) and Dataset B (on the right). While both templates covered the frontoparietal bilateral region, the optode template for Dataset A consisted of 8 sources (S) and 10 detectors (D), resulting in 18 channels. The optopde template for Dataset B consisted of 15 sources and 11 detectors, resulting in 32 channels.

#### 2.1.2 Dataset B

The dataset B consisted of 39 recordings collected from neurotypical primary school children (Mean age=12.00 years; SD=0.51) in South Africa. The data were recorded during an approximately 25-minute-long active numerical task requiring keypress responses. Although, unlike in Dataset A, movement was less of an issue, the main challenge was enough light penetration and absorption due to children’s predominantly darker skin and thicker hair. Data were acquired using the NIRScout system (NIRx Medical Technologies, LLC) with 15 sources and 11 detectors at wavelengths of 760 and 850. There were 32 channels located bilaterally in prefrontal and parietal cortices, sampled at 4.17 Hz (Fig. 1).

#### 2.1.3 Data preprocessing

All preprocessing steps were conducted using the NIRS Brain AnalyzIR Toolbox (Santosa et al., 2018), as one of the most commonly used software for fNIRS data (Yucel et al. 2025). For standardisation, both datasets were trimmed to include a maximum of 10 seconds before and after task onset and offset. The raw intensity values were then converted into optical density signals. After this, we first ran QT-NIRS on the uncorrected data (no MA correction) using the following generic parameters: QT=0.4, SCI=0.6, PSP=0.075, and a time window of 3 seconds. Next, we applied the motion artefact (MA) correction methods listed in Table 1 to the same datasets. After each correction, we again ran QT-NIRS with the same parameters. Finally, we extracted the resulting QT-NIRS metrics for the analysis, with the type of metrics differing between the simplified and advanced approaches (see Section 2.2 for details). Note that for Dataset A, the heartbeat frequency range in QT-NIRS was set to 0.5–2.5 Hz, corresponding to 30–150 beats per minute. For Dataset B, the upper limit was reduced to 2 Hz, because Dataset B had a lower sampling rate (4 Hz), and according to the Nyquist criterion, the maximum resolvable frequency is half the sampling rate (2 Hz). Retaining an upper limit of 2.5 Hz in this case would risk aliasing, where higher-frequency components are misrepresented in the signal. After this, we extracted the resulting QT-NIRS metrics for the analysis. The type of extracted metrics differed for the simplified and advanced approaches (See Section 2.2 for an in-depth description of the approaches). Our code for preprocessing is available at https://github.com/elizaveta-iva/methodological_study and https://osf.io/gf9sp/.

**Table 1.** Brief description of the MA correction methods used in our study.

| MA correction method | Description |
| --- | --- |
| Wavelet | Wavelet decomposes the signal into wavelet coefficients, sets those exceeding a statistical threshold (i.e., likely artefacts) to zero, and reconstructs a cleaned signal using the inverse transformation (Molavi & Dumont, 2012) |
| TDDR | TDDR down-weights large, infrequent fluctuations in the temporal derivative, assumed to result from motion, and reconstructs the signal by integrating the adjusted derivatives using a robust estimator (Fishburn et al., 2021). |
| Wavelet+TDDR | The combined use of both methods has been recommended, as they demonstrate comparable performance that is superior to other approaches. While TDDR effectively captures dynamic changes of artefacts in the time domain, Wavelet excels at high-resolution noise removal in the frequency domain (Guan et al., 2024). |
| Baseline PCA (bPCA) | Baseline PCA reduces global noise by removing principal components derived from a separate resting-state recording, assuming a consistent spatial structure of systemic signals across baseline and task data. We followed Franceschini et al. (2006) in removing components explaining up to 80% of spatial variance. |
| PCA | A task-only PCA approach uses the same logic as bPCA but derives components directly from the task data, which risks removing true brain signals if they share spatial patterns with systemic noise (Zhang et al., 2005). |

### 2.2 Approaches to using QT-NIRS for MA correction

#### 2.2.1 Preliminary heartbeat check

Since QT-NIRS relies on the presence of a detectable heartbeat in the signal, we first aimed to examine whether any of the MA correction methods eliminate this component. For this purpose, we synthetically generated a ground-truth dataset with a prominent cardiac pulsation and added motion artifacts, and fed it through all MA corrections algorithms at hand. The data was generated using the NIRS Brain AnalyzIR Toolbox’s default parameters (for physiological noise: cardiac amplitude = 0.25, respiration amplitude = 0.25, Mayer waves amplitude = 0.25; for motion artefact simulation: spikes per minute = 2, shifts per minute = 0.5), resulting in 30 recordings with 32 source–detector pairs (i.e., channels) and a signal length of 3001 seconds for each channel (Santosa et al., 2018). The raw synthesised data were then converted to optical density signals.

Power spectral density (PSD) analysis was performed on the optical density signals to assess preservation of physiological components after MA correction. For each channel, the dominant frequency within the 0.8-1.5 Hz cardiac band was identified. A cardiac peak was detected in 100% of channels across all six preprocessing pipelines. The mean peak frequency (1.04 Hz) across methods was stable, although for the PCA-based method, the mean peak showed a slight increase (mean = 1.10 Hz, SD = 0.21), possibly indicating distortion of the physiological signal. However, as the peak stayed within the typical range used for detecting the heartbeat in QT-NIRS (0.5 - 2.5 Hz), these results confirm that all MA correction methods retained the cardiac signal (Table 2).

**Table 2.**
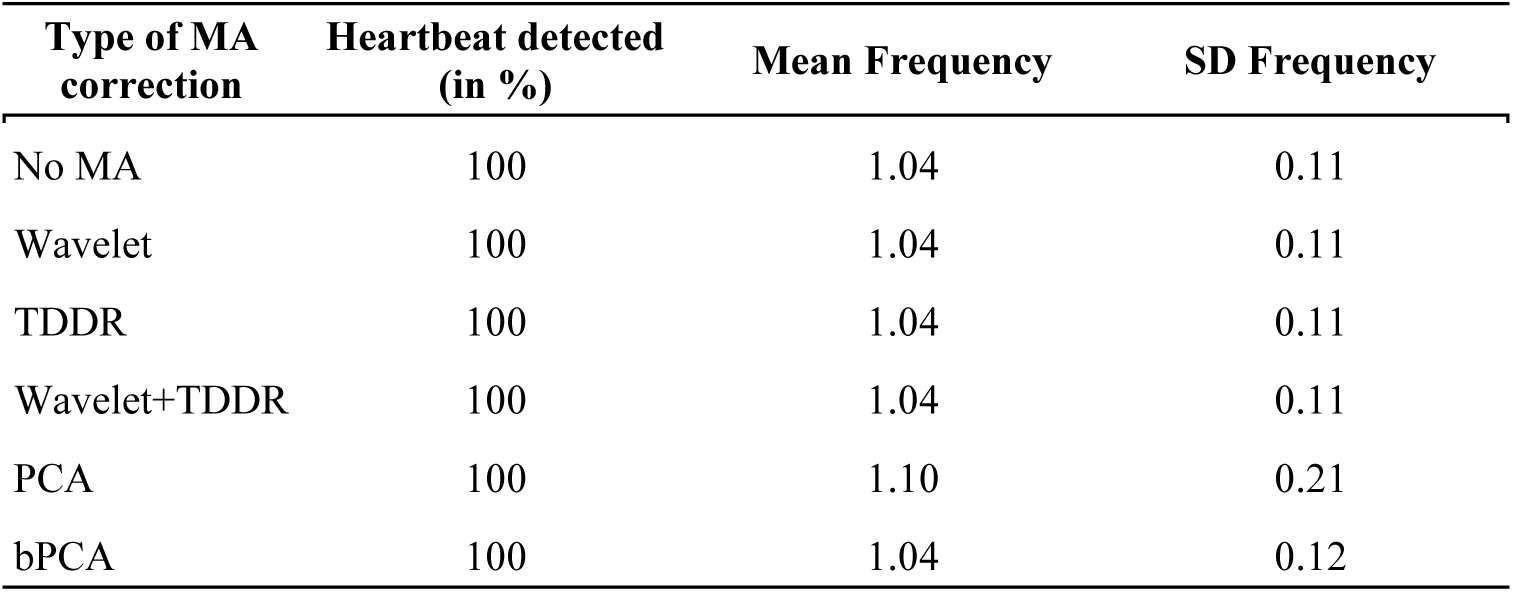
Heartbeat retention rates across MA correction methods.

| Type of MA correction | Heartbeat detected (in %) | Mean Frequency | SD Frequency |
| --- | --- | --- | --- |
| No MA | 100 | 1.04 | 0.11 |
| Wavelet | 100 | 1.04 | 0.11 |
| TDDR | 100 | 1.04 | 0.11 |
| Wavelet+TDDR | 100 | 1.04 | 0.11 |
| PCA | 100 | 1.10 | 0.21 |
| bPCA | 100 | 1.04 | 0.12 |

#### 2.2.2 Simplified approach

This approach requires no additional preparation or modification to QT-NIRS. The number of bad windows is simply extracted from the QT-NIRS reports after running them on the data with and without MA correction. In that case, the definition of the bad windows in QT_NIRS remains unchanged: a window is classified as bad if too few channels across such a time interval satisfy the predefined signal quality criteria. For example, in a case where we have 18 channels and set the QT at 0.4, a bad window would be the one in which fewer than 7 channels do not satisfy the PSP and SCI thresholds.

This definition implies a strong dependency of the number of bad windows on the QT value. For example, a QT value of 0.6 would mean that at least 60% of the data should have sufficient PSP and SCI values, and the QT value of only 0.2 is too liberal, suggesting the need for only 20% of the signal to meet our criteria. To see how much the results would be affected by the QT value, we additionally ran QT-NIRS with the QT setting at 0.2 and 0.6 in addition to 0.4, while keeping other settings the same (SCI=0.6, PSP=0.075, a time window of 3 seconds).

After running QT-NIRS on the data at the QT of 0.2, 0.4 and 0.6, we extracted the values from the *bad_windows* variable for each of the six MA correction methods (i.e., no MA, Wavelet, TDDR, Wavelet+TDDR, PCA, bPCA) for the subsequent analysis. Our code for data extraction is available at https://github.com/elizaveta-iva/methodological_study and https://osf.io/gf9sp/.

#### 2.2.3 Advanced approach

The advanced approach aimed at minimising the dependence of the bad windows calculation on other parameters, such as SCI and QT, and involved two key steps. First, we modified the original definition of bad windows in QT-NIRS in such a way that only time windows with clear evidence of artefacts were marked as bad:

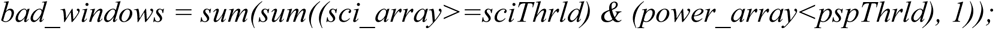

In this updated line, a window is considered bad when its PSP value is poor, but its SCI value is satisfactory. Note that the QT threshold is not important here anymore, which eliminates its potential influence (i.e., an arbitrary threshold of 0.2, 0.4, and 0.6 in our simplified approach) on the outcome. The good SCI here is critical, as the cases where SCI is poor are usually due to bad coupling between the optodes and the skin, which would distort fNIRS signals at both wavelengths to the point where heartbeat cannot be visually observed, making the data unreliable. When the signal passes the SCI threshold, it indicates that the fNIRS measurements are of sufficient quality to distinguish physiological components from noise. In this case, the PSP quantifies the strength of the motion artefact component, making it a valid measure of MA occurrence. Overall, the updated line eliminates the necessity to rely on QT and ensures that only cases that are truly MAs are defined as such.

We also introduced one more step to investigate the possibility that signal transformation from applying an MA correction method causes previously good (quality-wise) windows to be reclassified as bad windows, which would lead to false positives. To assess preservation of data quality post-correction, we calculated the proportion of good windows that remained unchanged before and after MA correction, stored in the *combo_array* value. We then calculated the efficiency score for each MA correction method using the following formula:

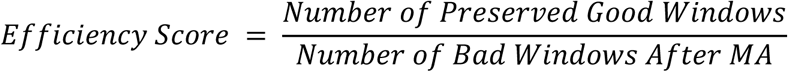

The efficiency score was calculated as the ratio between the number of good-quality windows preserved after MA correction and the number of bad-quality windows after MA. Higher values indicate better performance of the MA correction method. For the no MA condition, where the data was not corrected by any of the MA correction methods, we calculated the efficiency score by dividing the number of good windows by the number of bad windows.

Similarly to the simplified approach, the resulting number of bad windows and the efficiency scores were then used in the within-subject analysis. Our code is available at https://github.com/elizaveta-iva/methodological_study and https://osf.io/gf9sp/.

### 2.3 Comparison across MA methods

We analysed the effect of different MA correction methods on the number of bad windows using a non-parametric Friedman test. This test was chosen because the data violated the assumptions for normality and involved repeated measures from the same dataset across multiple MA correction methods. When the Friedman test indicated a significant main effect, post hoc comparisons were conducted to identify which specific correction methods differed from each other.

## 3 Results

### 3.1 Simplified approach

#### 3.1.1 Dataset A

At the QT of 0.4, a non-parametric Friedman test revealed a significant main effect of MA correction, χ²(5)=201.50, p<.001, Kendall’s W=0.67. Post hoc comparisons using Conover’s test with Benjamini-Hochberg correction revealed that the applications of TDDR and Wavelet+TDDR significantly decreased the number of bad windows compared to no MA correction, suggesting improvement in the signal quality. TDDR did not differ significantly from Wavelet+TDDR. The application of Wavelet did not significantly change the number of bad windows. Surprisingly, the application of PCA and bPCA significantly increased the number of bad windows, which meant worsening the signal quality. Out of these two, PCA worsened the signal quality significantly more than bPCA. Further comparisons revealed that the application of Wavelet+TDDR, as well as TDDR, significantly decreased the number of bad windows compared to Wavelet. Compared to both PCA and bPCA, Wavelet, TDDR, and Wavelet+TDDR performed significantly better. To summarise, at the QT of 0.4, TDDR and Wavelet+TDDR significantly reduced the number of bad windows, whereas PCA and bPCA significantly increased them, with PCA performing worse than bPCA.

We additionally tested the change in the outcome when QT is decreased to 0.2 and increased to 0.6 due to its arbitrary nature in the calculation of the number of bad windows. At the QT of 0.2, a non-parametric Friedman test revealed a significant main effect of MA correction, χ²(5)=196.32, p<.001, Kendall’s W=0.65. Post hoc comparisons using Conover’s test with Benjamini-Hochberg correction revealed that the application of Wavelet, TDDR and Wavelet+TDDR significantly decreased the number of bad windows compared to no MA correction, suggesting improvement in the signal quality. The difference between Wavelet and Wavelet+TDDR was not significant, as well as between Wavelet and TDDR. However, TDDR was significantly worse compared to Wavelet+TDDR. Surprisingly, the application of PCA and bPCA significantly increased the number of bad windows, which meant worsening the signal quality. Out of them, PCA was significantly worse than bPCA. To summarise, at the QT of 0.2, Wavelet, TDDR, and Wavelet+TDDR significantly reduced the number of bad windows, whereas PCA and bPCA significantly increased them, with PCA performing worse than bPCA.

At the QT of 0.6, a non-parametric Friedman test revealed a significant main effect of MA correction, χ²(5)=189.67, p<.001, Kendall’s W=0.63. Post hoc comparisons using Conover’s test with Benjamini-Hochberg correction revealed that the application of Wavelet, Wavelet+TDDR, PCA and bPCA significantly increased the number of bad windows compared to no MA correction, which meant worsening the signal quality. There was no difference between Wavelet and Wavelet+TDDR, although both Wavelet and Wavelet+TDDR were still significantly better than PCA and bPCA. The application of PCA led to significantly more bad windows than that of bPCA. At the same time, the application of the TDDR was not significant. Overall, at the QT of 0.6, no MA correction method significantly decreased the number of bad windows; on the contrary, application of all tested MA correction methods except TDDR led to an increase in the number of bad windows.

Across QT thresholds of 0.2 and 0.4, TDDR and Wavelet+TDDR consistently showed the most improvement in reducing the number of bad windows. PCA and bPCA consistently worsened signal quality, with PCA performing the worst in all cases. At QT=0.6, none of the tested MA correction methods significantly reduced the number of bad windows, and all except TDDR led to an increase in the number of bad windows (see Table 3, Fig. 2 and Fig. 3).

**Fig. 2.**
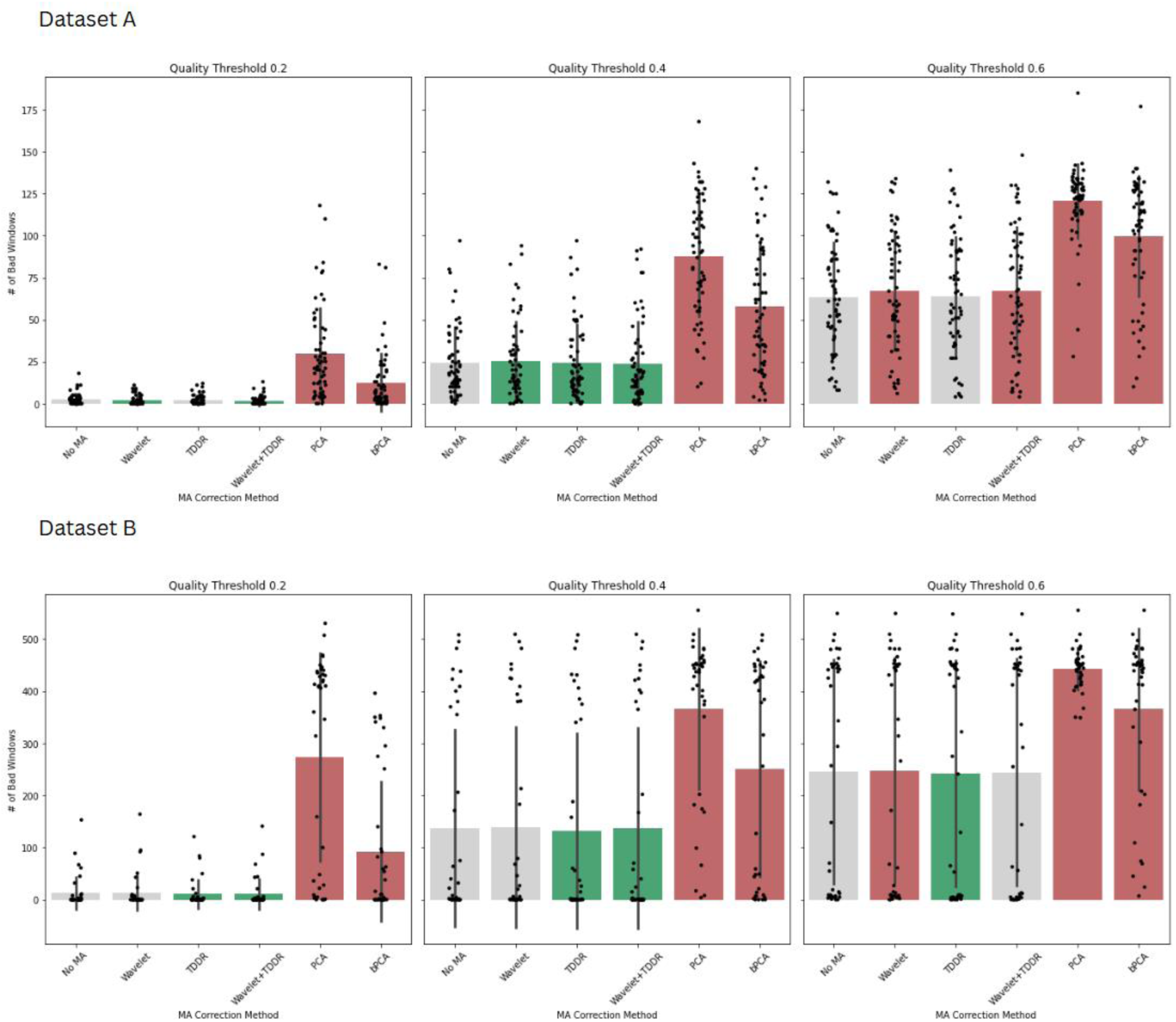
Number of bad windows remaining after MA correction across datasets, quality thresholds, and correction methods after using the simplified approach. Violin plots show the distribution of bad windows remaining after applying each MA correction method across two datasets (Dataset A: top row; Dataset B: bottom row) and three quality thresholds (QT of 0.2, 0.4, 0.6; left to right columns). MA methods that significantly reduced the number of bad windows compared to no correction are outlined in green, while those that significantly increased the number of bad windows are outlined in red. Significant post hoc comparisons between correction methods (excluding those that significantly worsened data quality) are indicated with brackets and p-values.

**Fig. 3.**
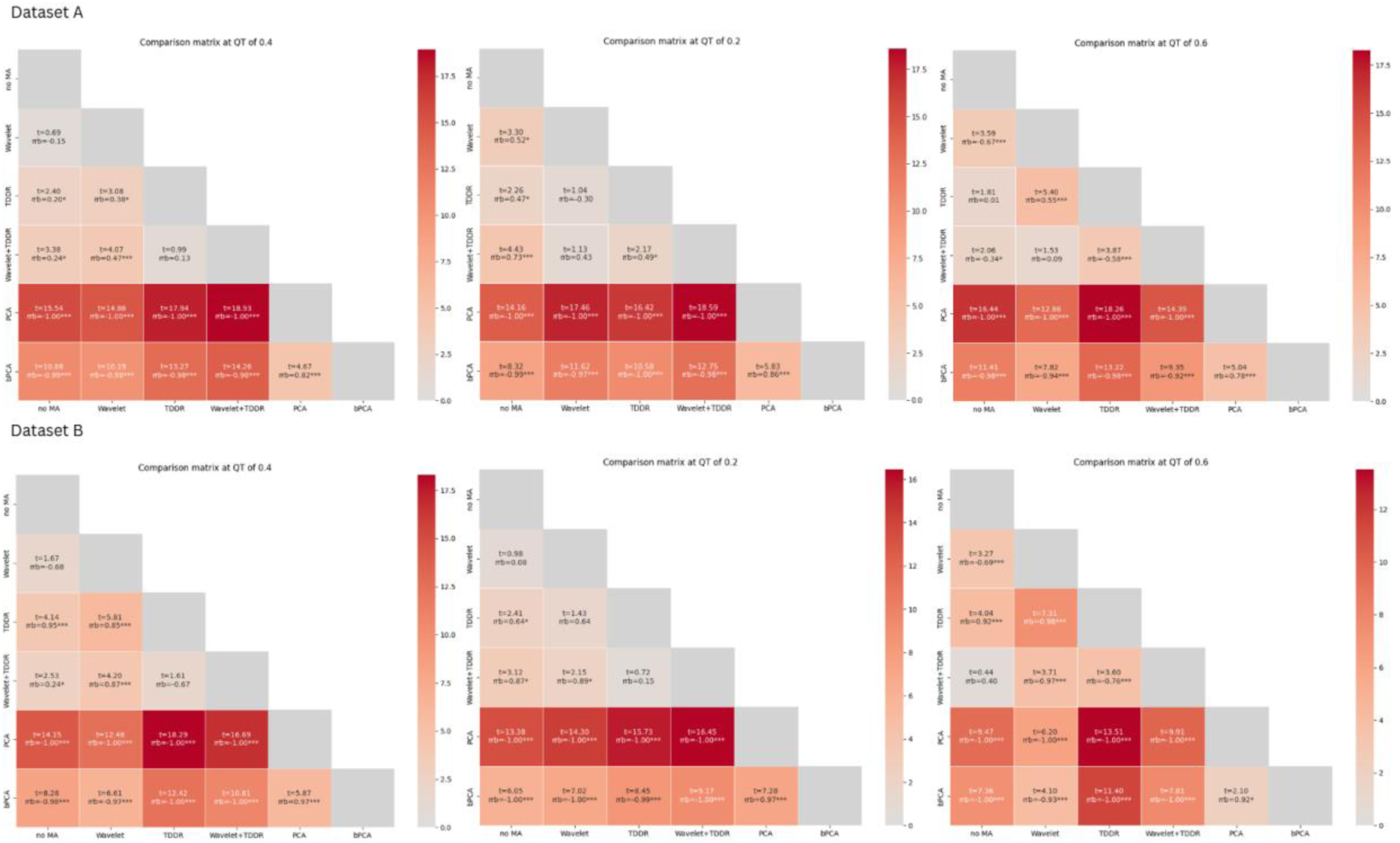
Pairwise comparisons of motion artefact (MA) correction methods in two datasets. Each matrix shows the results of post hoc comparisons between MA methods (Dataset A: top matrix; Dataset B: bottom matrix). Values are read from column to row and report the t statistic (t), significance level (p), and effect size (rrb). Positive rrb values indicate that the method in the column outperformed the method in the row, whereas negative values indicate the opposite. Significant differences are marked with an * for p<.05, ** for p<.01 and *** for p<.001.

**Table 3.** Descriptive statistics of the remaining number of bad windows and their percentage from the total number of windows after using the simplified approach across two datasets (A and B) and three quality thresholds (QT=0.2, 0.4, 0.6). Lower values reflect more effective artefact removal.

| Type of MA correction | QT 0.2 |  | QT 0.4 |  | QT 0.6 |  |
| --- | --- | --- | --- | --- | --- | --- |
|  | M (%) | SD (%) | M (%) | SD (%) | M (%) | SD (%) |
| **Dataset A (average N of windows = 133.65, SD = 9.11)** |  |  |  |  |  |  |
| **No MA** | 2.62 (1.95%) | 3.43 (2.55%) | 24.48 (18.27%) | 21.08 (15.69%) | 63.25 (47.24%) | 33.03 (24.38%) |
| **Wavelet** | 1.82 (1.34%) | 2.76 (2.01%) | 25.5 (18.95%) | 23.30 (17.16%) | 67.42 (50.26%) | 35.47 (25.92%) |
| **TDDR** | 2.07 (1.54%) | 2.93 (2.19%) | 24.13 (17.95%) | 23.12 (17.08%) | 63.92 (47.66%) | 35.65 (25.97%) |
| **Wavelet+TDDR** | 1.53 (1.12%) | 2.59 (1.85%) | 23.97 (17.73%) | 24.70 (17.94%) | 67.08 (49.98%) | 38.11 (27.75%) |
| **PCA** | 29.88 (22.14%) | 27.43 (19.90%) | 87.92 (65.58%) | 36.06 (25.80%) | 120.62 (90.21%) | 22.18 (15.09%) |
| **bPCA** | 12.58 (9.26%) | 17.11 (12.34%) | 57.7 (42.81%) | 38.39 (27.59%) | 99.63 (74.30%) | 35.97 (25.80%) |
| **Dataset B (average N of windows = 456.28, SD = 29.41)** |  |  |  |  |  |  |
| **No MA** | 12.46 (2.63%) | 31.13 (6.42%) | 136.74 (29.65%) | 191.36 (41.13%) | 245.59 (53.08%) | 217.88 (46.58%) |
| **Wavelet** | 12.87 (2.70%) | 33.70 (6.90%) | 139.64 (30.30%) | 194.74 (41.94%) | 248.13 (53.66%) | 218.18 (46.66%) |
| **TDDR** | 10.87 (2.30%) | 27.19 (5.64%) | 132.49 (28.71%) | 189.11 (40.59%) | 241.92 (52.26%) | 219.03 (46.84%) |
| **Wavelet+TDDR** | 10.87 (2.28%) | 28.92 (5.94%) | 137.08 (29.73%) | 194.19 (41.80%) | 244.59 (52.86%) | 219.62 (46.99%) |
| **PCA** | 272.92 (59.35%) | 201.49 (43.45%) | 366.44 (80.22%) | 155.66 (33.00%) | 442.90 (97.16%) | 40.42 (7.21%) |
| **bPCA** | 92.51 (20.15%) | 135.58 (29.56%) | 250.77 (54.66%) | 208.26 (45.18%) | 365.72 (79.97%) | 156.70 (33.30%) |

#### 3.1.2 Dataset B

At the QT of 0.4, a non-parametric Friedman test revealed a significant main effect of MA correction, χ²(5)=140.44, p<.001, Kendall’s W=0.72. Post hoc comparisons using Conover’s test with Benjamini-Hochberg correction revealed that application of the TDDR and Wavelet+TDDR significantly reduced the number of bad windows when compared to no MA correction. The application of Wavelet did not significantly affect the number of bad windows. The number of bad windows after TDDR did not significantly differ from Wavelet+TDDR. Additionally, the number of bad windows was significantly increased when PCA and bPCA were applied. After applying PCA, the data had significantly more bad windows than bPCA. Overall, at the QT of 0.4, TDDR and Wavelet+TDDR improved signal quality, Wavelet had no significant effect, and PCA-based methods worsened it, with PCA performing the worst.

At the QT of 0.2, a non-parametric Friedman test revealed a significant main effect of MA correction, χ²(5)=132.85, p<.001, Kendall’s W=0.68. Post hoc comparisons using Conover’s test with Benjamini-Hochberg correction revealed that only the application of Wavelet+TDDR and TDDR significantly reduced the number of bad windows, when compared to no MA correction, although there was no difference between them. There was also no difference when the Wavelet was applied. Additionally, the number of bad windows was significantly increased when PCA and bPCA were applied. Out of those two, PCA had a significantly worse effect on the number of bad windows. Overall, at QT=0.2, only TDDR and Wavelet+TDDR improved signal quality; Wavelet alone had no significant effect, and PCA-based methods worsened it, with PCA again performing the worst.

When the QT was set at 0.6, a non-parametric Friedman test revealed a significant main effect of MA correction, χ²(5)=112.81, p<.001, Kendall’s W=0.58. Post hoc comparisons using Conover’s test with Benjamini-Hochberg correction revealed that, compared to no MA correction, only TDDR significantly reduced the number of bad windows. Wavelet+TDDR did not have a significant effect on the number of bad windows. At the same time, Wavelet, PCA and bPCA had a significantly higher number of bad windows. Out of these, Wavelet was still significantly better than PCA and bPCA, meaning its effect on the data quality was not as detrimental as in the case of PCA and bPCA. Similarly to previous results, PCA was significantly worse than bPCA. To summarise, at QT of 0.6, application of TDDR decreased the number of bad windows, application of Wavelet+TDDR did not lead to any changes in the data quality, and the application of Wavelet, PCA, and bPCA worsened it.

Similar to the outcome for Dataset A, at QT thresholds of 0.2 and 0.4, TDDR and Wavelet+TDDR consistently showed the most improvement in reducing the number of bad windows. PCA and bPCA consistently worsened signal quality, with PCA performing the worst in all cases. At QT=0.6, only TDDR significantly reduced the number of bad windows; the application of Wavelet+TDDR did not lead to any changes, while Wavelet, PCA and bPCA significantly worsened the quality of data (see Table 3, Fig. 2 and Fig. 3).

### 3.2 Computationally advanced approach

In the advanced approach, we analysed the effect of different MA correction methods on both the number of bad windows and the efficiency scores using a non-parametric Friedman test. Similarly to the analysis of the simplified approach, this test was chosen because the data violated the assumptions of normality and involved repeated measures from the same dataset across multiple correction methods. When the Friedman test indicated a significant main effect, post hoc comparisons were conducted to identify which specific correction methods differed from each other. See Table 4, Fig. 4 and Fig. 5 for visualised findings.

**Fig. 4.**
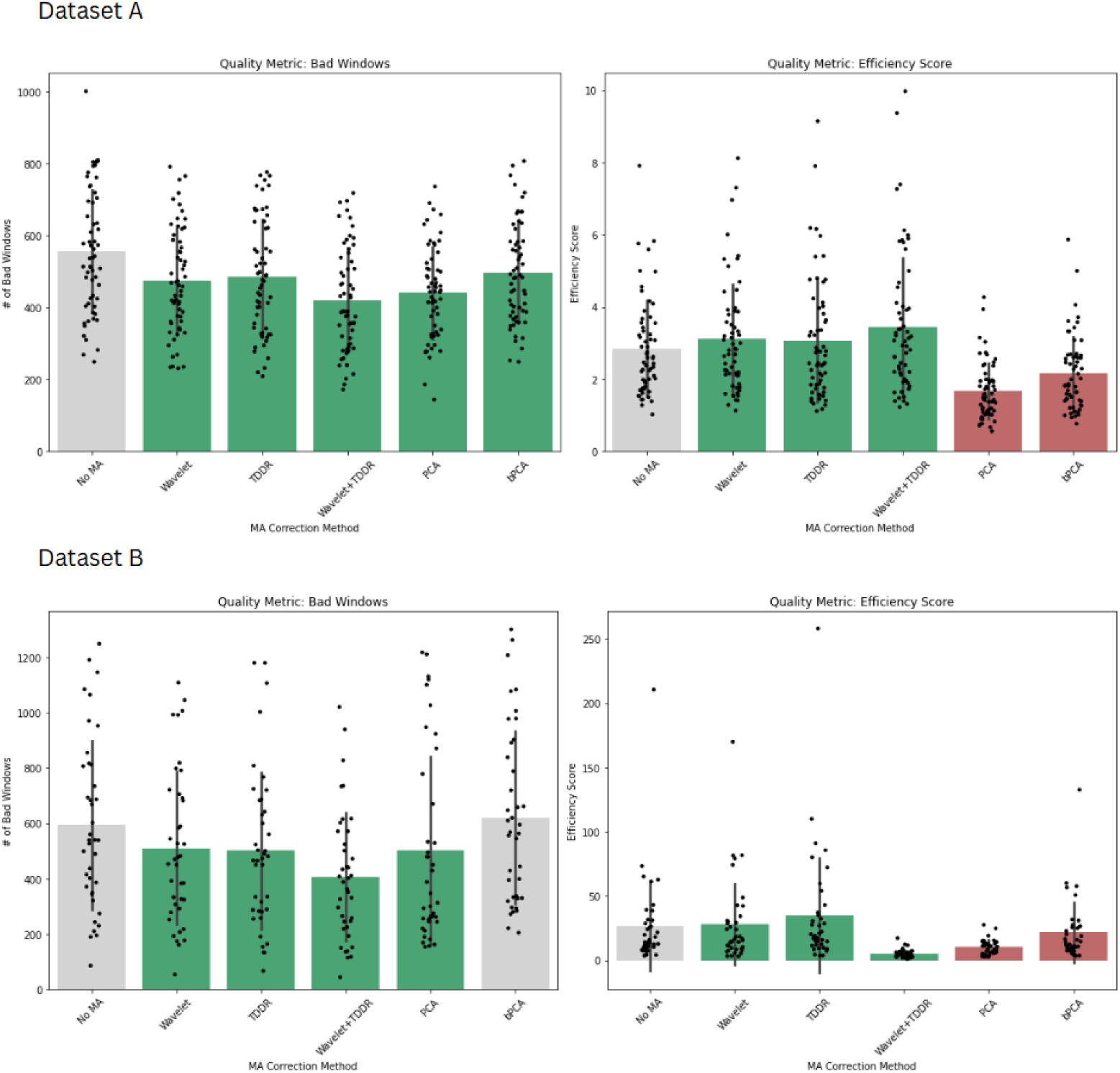
Number of bad windows and efficiency scores after MA correction across datasets after using the advanced approach. Violin plots show the distribution of bad windows remaining after applying each MA correction method across two datasets (Dataset A: top row; Dataset B: bottom row) and three quality thresholds (QT of 0.2, 0.4, 0.6; left to right columns). MA methods that significantly reduced the number of bad windows compared to no correction are outlined in green, while those that significantly increased the number of bad windows are outlined in red. Significant post hoc comparisons between correction methods (excluding those that significantly worsened data quality) are indicated with brackets and p-values.

**Fig. 5.**
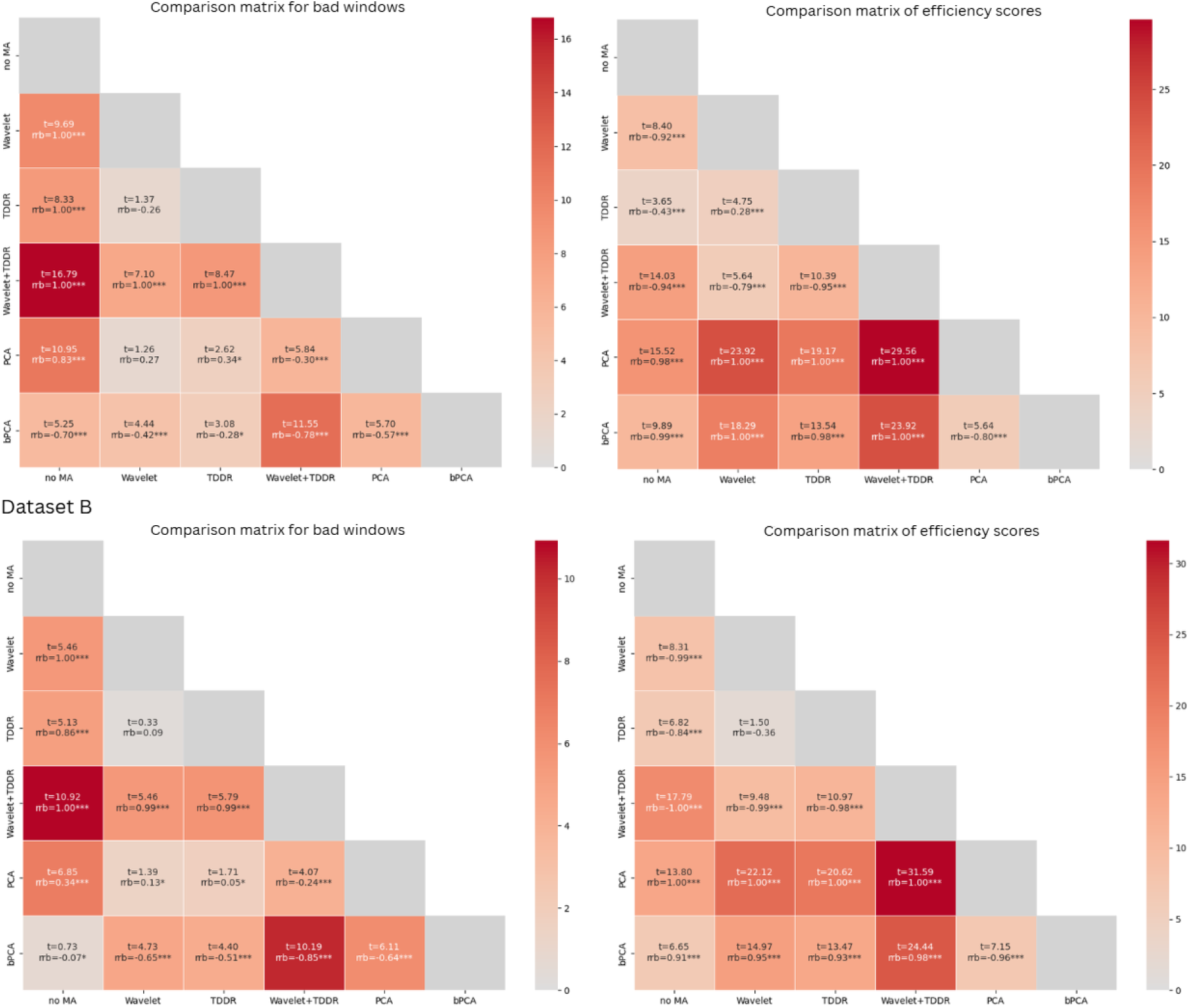
Pairwise post hoc comparisons between MA correction methods (Dataset A: top matrix; Dataset B: bottom matrix). Values are read from column to row and include the t statistic (t), significance level (p), and effect size (rrb). Positive rrb values indicate that the method in the column outperformed the method in the row, whereas negative values indicate the opposite. Separate matrices are shown for bad windows and efficiency scores. Significant differences are marked with an * for p<.05, ** for p<.01 and *** for p<.001.

**Table 4.** Descriptive statistics of the absolute number of bad windows, their percentage from the total number of windows, and efficiency scores after using the advanced approach across two datasets (A and B). Lower numbers of bad windows and higher efficiency scores suggest more effective MA removal.

| Type of MA correction | Number of bad windows |  | Efficiency scores |  |
| --- | --- | --- | --- | --- |
|  | M (%) | SD (%) | M | SD |
| **Dataset A (average N of windows = 2405.70, SD = 164.07)** |  |  |  |  |
| **No MA** | 557.95<br>(23.14%) | 170.87<br>(6.68%) | 2.85 | 1.34 |
| **Wavelet** | 474.00<br>(19.67%) | 146.21<br>(5.76%) | 3.10 | 1.54 |
| **TDDR** | 485.45<br>(20.16%) | 160.97<br>(6.49%) | 3.05 | 1.69 |
| **Wavelet+TDDR** | 421.27<br>(17.50%) | 143.81<br>(5.79%) | 3.44 | 1.94 |
| **PCA** | 441.08<br>(18.36%) | 129.01<br>(5.29%) | 1.68 | 0.77 |
| **bPCA** | 495.88<br>(20.65%) | 140.98<br>(5.86%) | 2.15 | 1.03 |
| **Dataset B (average N of windows = 14601.03, SD = 941.18)** |  |  |  |  |
| **No MA** | 593.69<br>(4.10%) | 307.05<br>(2.19%) | 21.68 | 23.64 |
| **Wavelet** | 510.36<br>(3.52%) | 279.31<br>(2.00%) | 26.73 | 35.19 |
| **TDDR** | 501.77<br>(3.46%) | 286.10<br>(2.02%) | 28.05 | 32.03 |
| **Wavelet+TDDR** | 405.77<br>(2.80%) | 233.62<br>(1.65%) | 35.24 | 45.12 |
| **PCA** | 500.82<br>(3.47%) | 344.56<br>(2.44%) | 5.29 | 3.25 |
| **bPCA** | 620.95<br>(4.28%) | 315.70<br>(2.27%) | 10.19 | 5.75 |

#### 3.2.1 Dataset A

A non-parametric Friedman test between the number of bad windows after preprocessing revealed a significant main effect of the MA correction, χ²(5)=155.66, p<.001, Kendall’s W=0.52. Post-hoc tests using Conover’s test with Benjamini-Hochberg correction revealed that while all five MA correction methods significantly lowered number of bad windows when compared to no MA correction, application of the Wavelet+TDDR had significantly reduced the number of bad windows than Wavelet, TDDR, bPCA, and PCA. Therefore, although all MA correction methods performed well, Wavelet+TDDR had the least number of bad windows as compared to the other MA correction methods.

When compared by the efficiency scores, a non-parametric Friedman test between the efficiency scores after preprocessing revealed a significant main effect of the MA correction, χ²(5)=242.30, p<.001, Kendall’s W=0.81. Post hoc comparisons using Conover’s test with Benjamini-Hochberg correction revealed that the dataset preprocessed with Wavelet, TDDR, and Wavelet+TDDR had higher efficiency scores compared to no MA correction. Among them, Wavelet+TDDR had a significantly higher efficiency score than both Wavelet and TDDR. At the same time, PCA and bPCA had significantly lower efficiency scores than when no MA correction was applied. Overall, Wavelet+TDDR was the most efficient method, while the application of PCA and bPCA methods led to significant distortion of the data.

#### 3.2.2 Dataset B

A non-parametric Friedman test between the number of bad windows after applying five MA correction methods and control revealed a significant main effect of MA correction, χ²(5)=242.30, p<.001, Kendall’s W=0.68. The application of all MA correction methods except bPCA significantly decreased the number of bad windows, compared to no MA condition. Post hoc comparisons using Conover’s test with Benjamini-Hochberg correction revealed that out of the MA correction methods that significantly decreased the number of bad windows, data corrected with Wavelet+TDDR had the least number of bad windows than with Wavelet, TDDR and PCA. At the same time, the difference between data preprocessed with Wavelet and TDDR was not significant. To summarise, although all MA correction methods except bPCA decreased the number of bad windows, Wavelet+TDDR did significantly better than the others. When compared by the efficiency scores, a non-parametric Friedman test between the efficiency scores when MA correction methods were applied and no MA correction revealed a significant main effect of MA correction, χ²(5)=169.83, p<.001, Kendall’s W=0.87. Post hoc comparisons using Conover’s test with Benjamini-Hochberg correction revealed that Wavelet, TDDR and Wavelet+TDDR had higher efficiency scores than no MA. Among them, Wavelet+TDDR had a significantly higher efficiency score than both Wavelet and TDDR. Wavelet and TDDR did not differ. At the same time, PCA and bPCA had significantly lower efficiency scores than no MA. Between them, PCA had a significantly lower score. To summarise, the analysis of the efficiency scores revealed that Wavelet+TDDR had the highest efficiency score, meaning that it improved bad windows while not distorting good windows best. The application of bPCA and especially PCA led to significant decrease in the efficiency scores, meaning that they distorted more good quality windows than improved bad quality ones.

## 4 Discussion

This paper aimed to contribute to the development of shared standards in the preprocessing and reporting of fNIRS data (Kelsey et al., 2023; Schroeder et al., 2023; Yücel et al., 2021; Yücel et al., 2025). We proposed a QT-NIRS-based algorithm in simplified and advanced approaches for selecting the optimal MA correction method per dataset. Our proposed approach is reproducible, replicable, and accessible to researchers with varying levels of programming expertise, and effective across a wide range of datasets. While we have successfully demonstrated our algorithm in the results section, below, we discuss some considerations for both simplified and advanced approaches.

### 4.1 Simplified approach

At the generic setting for the QT-NIRS (QT=0.4, SCI=0.6, PSP=0.075, a time window of 3 seconds), application of either Wavelet+TDDR or TDDR decreased the most bad windows for Datasets A and B. The results remained similar when the QT was decreased to 0.2, except that in Dataset A, application of Wavelet also led to data quality improvement. However, when QT was increased to 0.6, a statistically significant improvement was only observed in Dataset B, when TDDR was applied. Surprisingly, the application of Wavelet and Wavelet+TDDR worsened the data quality in Dataset A, and Wavelet worsened the data quality in Dataset B. Nonetheless, the effect of Wavelet and Wavelet+TDDR was weak in either dataset.

Comparatively increased conservativeness of the QT of 0.6, alongside the approach to calculating the number of bad windows, could explain the surprising change in results. In the simplified approach, bad windows are identified indirectly, with SCI, PSP, and the QT used together as a proxy. As such, adopting a more liberal QT may reduce the influence of other parameters (e.g., SCI) on the final output. Given that in both datasets the effect from applying Wavelet, TDDR, or Wavelet+TDDR only drastically changed at QT of 0.6, as well as given their small effect sizes at that measure, we suggest that QT of 0.6 might be less stable compared to when QT is set to 0.2 or 0.4.

As such, it seems fair to suggest that based on the more agreeable results at QT of 0.2 and 0.4, the application of Wavelet+TDDR and TDDR was overall beneficial for the datasets and confirms previous research suggestions about the high effectiveness of both Wavelet and TDDR (Guan et al., 2024). While we do not recommend using a low QT value of 0.2 for channel pruning, we advise that users intending to apply the simplistic approach run QT-NIRS separately: once for channel pruning and once for selecting the MA correction method. This way, channel pruning would still be able to rely on the conservative QTs, ensuring better quality of data. At the same time, running QT-NIRS for the second time with more liberal settings would decrease the influence of SCI when deciding on the MA correction method. Overall, the simplistic approach should be applied with much greater caution than the advanced approach.

Quite surprisingly, in both datasets, we observed a substantial increase in the number of bad windows when PCA and bPCA were used, regardless of the QT threshold. We suggest that one should be more cautious with using those MA correction methods, as the results of their application are quite alarming. This may be because PCA-based methods can remove not only noise but also meaningful signal variance, leading to increased instability. Their reliance on variance decomposition might also amplify small fluctuations rather than suppress them. This supports the previous research on comparing the MA correction methods that highlighted instability of the PCA-based methods and their lower performance, compared to Wavelet or TDDR (Brigadoi et al., 2014; Guan et al., 2024). Overall, these results suggest that Wavelet and TDDR are more reliable choices for MA correction.

Overall, the results also suggest that, despite QT value playing a definitive role in calculating the number of bad windows, the changes to the QT at 0.2 and 0.4 did not considerably destabilise the outcome. PCA and bPCA underperformed under all tested conditions, and the top-performing MA correcting method was a consistent combination of TDDR or Wavelet+TDDR, and sometimes Wavelet, all three of which did not seem to have strong differences (as observed in Figure X and Table X) in any case. That said, if one chooses to rely on the simplified approach for selecting the most appropriate MA correction method, it would be wise to see how the results change under different QT values, to ensure better stability of the choice. Given the potential instability of QT of 0.6, we recommend running QT-NIRS at a lower QT setting when using QT-NIRS for choosing the best MA correction methods.

### 4.2 Computationally advanced approach

Similarly to the simplified approach, in both datasets, the combination of Wavelet+TDDR was found to be the most successful, as evidenced by both the reduced number of bad windows and the efficiency scores. These much more definitive results suggest that at least within the MA correction methods tested, Wavelet+TDDR was the most beneficial for the data, as recently proposed by Guan et al. (2024).

Introducing an additional check by calculating the efficiency scores also helped recognise whether the MA correction methods distorted the good quality segments of the data. Especially in Dataset A, based on the number of bad windows pre-and post-MA correction, PCA and bPCA showed promising results. However, when they were compared by their efficiency scores, they showed a worse result than a condition with no MA correction by damaging the good quality segments of the data far more often than they improved the bad quality segments. This result was in line with the conclusions made in the previous studies (Brigadoi et al., 2014), as well as the results from the simplified approach that did not have the efficiency scores calculated to keep the approach as easy to conduct as possible. Overall, adding the efficiency scores to the advanced approach seems to be a helpful solution for minimising bias in the results.

### 4.3 General recommendations

Regardless of whether a computationally advanced or simplified approach is preferred, it is crucial to consider the influence of parameter settings on the outcome. Each dataset is unique and may require further adjustment of QT-NIRS parameters, which are oftentimes defined empirically (Hernandez & Pollonini, 2020). For example, in participants with lower heart rates, such as athletes, increasing the window length to 4 or 5 seconds may help better capture the heartbeat signal. Or as in our case, knowing that your participants can be quite active and have an increased number of MA can prompt one to have a more liberal threshold for SCI. Likewise, values for PSP, SCI, and the overall QT can be modified accordingly, and we encourage everyone to adjust them for their study purposes. As changes to any of these parameters can significantly affect the results, we strongly recommend that all parameter values be reported in the papers to ensure transparency and reproducibility (Kelsey et al., 2023).

## 3 Conclusion

The proposed algorithm provides a reproducible and transparent framework for selecting an appropriate MA correction method applicable across various datasets. Although the computations were slightly different for the simplified and advanced approaches, they yielded similar recommendations in this order: Wavelet, TDDR or Wavelet+TDDR improved the signals, while PCA and bPCA worsened the signals. The choice between which approach to use is a matter of preference and computational skills. The advanced approach utilises more complicated analysis, but returns more straightforward and robust results. The simplified approach, on the other hand, is much easier to apply, but the results become slightly less stable as they depend on confounding variables. Regardless of using each of these approaches, they will significantly help to make an informed choice on which MA correction method is the most suitable for a dataset.

## Acknowledgments

We are grateful to Kathleen Fonseca (Department of Childhood Education, University of Johannesburg, South Africa) for collecting and providing the Dataset B.

This work was funded by the Faculty of Health and Medical Sciences at the University of Surrey to EI (TF3129), the FRSF grant from the Faculty of Health and Medical Sciences at the University of Surrey to MS (AC0643) and the National Research Foundation South Africa Research Chair to MS (Grant number: SARC98573).

## Data Availability Statement

Upon publication, the data and analytic code necessary to reproduce the analyses presented here will be publicly accessible and available at https://github.com/elizaveta-iva/methodological_study and https://osf.io/gf9sp/.

